# Further humoral immunity evasion of emerging SARS-CoV-2 BA.4 and BA.5 subvariants

**DOI:** 10.1101/2022.08.09.503384

**Authors:** Fanchong Jian, Yuanling Yu, Weiliang Song, Ayijiang Yisimayi, Lingling Yu, Yuxue Gao, Na Zhang, Yao Wang, Fei Shao, Xiaohua Hao, Yanli Xu, Ronghua Jin, Youchun Wang, Xiaoliang Sunney Xie, Yunlong Cao

**Affiliations:** Biomedical Pioneering Innovation Center (BIOPIC), Peking University, Beijing, P.R. China; College of Chemistry and Molecular Engineering, Peking University, Beijing, P.R. China; Division of HIV/AIDS and Sex-transmitted Virus Vaccines, Institute for Biological Product Control, National Institutes for Food and Drug Control (NIFDC), Beijing, P.R. China; School of Life Sciences, Peking University, Beijing, P.R. China; Changping Laboratory, Beijing, P.R. China; Joint Graduate Program of Peking-Tsinghua-NIBS, Academy for Advanced Interdisciplinary Studies, Peking University, Beijing, China; Beijing Ditan Hospital, Capital Medical University, Beijing, P.R. China

## Abstract

Multiple BA.4 and BA.5 subvariants with R346 mutations on the spike glycoprotein have been identified in various countries, such as BA.4.6/BF.7 harboring R346T, BA.4.7 harboring R346S, and BA.5.9 harboring R346I. These subvariants, especially BA.4.6, exhibit substantial growth advantages compared to BA.4/BA.5. In this study, we showed that BA.4.6, BA.4.7, and BA.5.9 displayed higher humoral immunity evasion capability than BA.4/BA.5, causing 1.5 to 1.9-fold decrease in NT50 of the plasma from BA.1 and BA.2 breakthrough-infection convalescents compared to BA.4/BA.5. Importantly, plasma from BA.5 breakthrough-infection convalescents also exhibits significant neutralization activity decrease against BA.4.6, BA.4.7, and BA.5.9 than BA.4/BA.5, showing on average 2.4 to 2.6-fold decrease in NT50. For neutralizing antibody drugs, Bebtelovimab remains potent, while Evusheld is completely escaped by these subvariants. Together, our results rationalize the prevailing advantages of the R346 mutated BA.4/BA.5 subvariants and urge the close monitoring of these mutants, which could lead to the next wave of the pandemic.

## Main

Severe acute respiratory syndrome coronavirus 2 (SARS-CoV-2) BA.4 and BA.5 lineages have been the dominant strain in most regions around the globe and are continuously gaining additional mutations on the RBD ^1–3^. Multiple BA.4 and BA.5 subvariants with R346 mutations on the spike glycoprotein have been identified in various countries, such as BA.4.6/BF.7 harboring R346T, BA.4.7 harboring R346S, and BA.5.9 harboring R346I (Fig. S1). These subvariants, especially BA.4.6, exhibit growth advantages compared to other variants, including original BA.4/BA.5 ^4^. Previous studies have identified R346 as an important immunogenic residue, and R346 mutations would escape a large group of neutralizing antibodies (NAbs) ^3^. Unlike R346K carried by BA.1.1, which did not cause significant immune evasion, mutations from Arg to Thr/Ser/Ile correspond to a stronger shift in chemical properties ^5,6^. Therefore, the efficacy of vaccines and NAb drugs against these BA.4/BA.5 sublineages needs immediate evaluation.

In this study, we measured the neutralizing titers of plasma samples against the SARS-CoV-2 BA.4/BA.5 subvariants with R346 mutations. The plasma samples are obtained from vaccinated individuals without SARS-CoV-2 infection or with BA.1/BA.2/BA.5 breakthrough infection (Table S1). Vesicular stomatitis virus (VSV)-based pseudoviruses were used in the neutralization assays.

Plasma samples from individuals who received triple doses of CoronaVac without infection showed 1.5 to 1.7-fold decrease in 50% neutralization titers (NT50) against BA.4/BA.5 with R346I (BA.5.9), R346T (BA.4.6 and BF.7), and R346S (BA.4.7), in comparison to the NT50 against BA.4/BA.5 (Fig. 1A). Similar reduction in neutralization titers was also observed in plasma from BA.1 or BA.2 breakthrough infection convalescents (Fig. 1B-C). Importantly, BA.4/BA.5 with R346I/T/S could significantly evade neutralization by plasma samples from BA.5 breakthrough infection, exhibiting a 2.4 to 2.6-fold decrease in NT50 (Fig. 1D). These results indicate the strong humoral immunity evasion capability of BA.4/BA.5 sublineages with R346 mutations, suggesting these sublineages, including BA.4.6, BA.4.7, BA.5.9, and BF.7, would gain an advantage in transmissibility under the global background of the pandemic caused by BA.4/BA.5.

**Figure 1.**
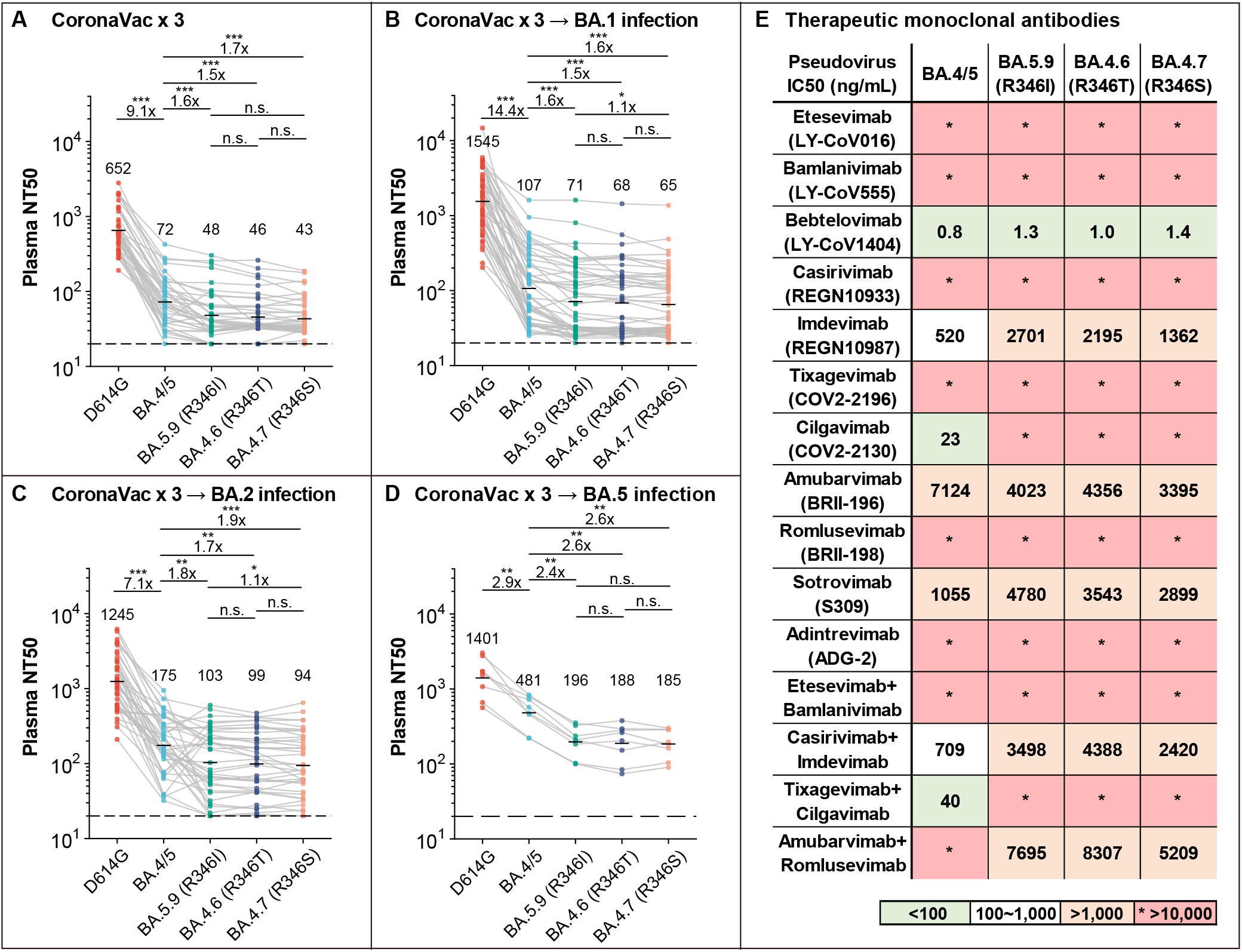
Efficacy of convalescent plasma and neutralizing antibody drugs against BA.4/BA.5 subvariants with mutations on spike R346. (A-D) NT50 against SARS-CoV-2 D614G, BA.4/5, BA.5.9 (BA.4/5+R346I), BA.4.6 (BA.4/5+R346T), BA.4.7 (BA.4/5+R346S) pseudovirus by plasma samples from individuals who received 3 doses CoronaVac (A, N=40), and those who received 3 doses CoronaVac followed by BA.1 breakthrough infection (B, N=50), BA.2 breakthrough infection (C, N=39), or BA.5 breakthrough infection (D, N=8). Geometric mean titers (GMT) are annotated above each group. *p < 0.05; **p < 0.01; ***p<0.001. P-values are calculated using a two-tailed Wilcoxon sign-rank test of paired samples. (E) Neutralizing activities against SARS-CoV-2 D614G and BA.4/5 subvariants pseudovirus of therapeutic neutralizing antibodies. Background colors indicate neutralization levels. Green, IC50 < 100 ng/mL; white, 100 ng/mL < IC50 < 1,000 ng/mL; red, IC50 > 1,000 ng/mL. *, IC50 > 10, 000 ng/mL.

Next, we evaluated the pseudovirus-neutralizing activities of the approved neutralizing antibody drugs, including 11 monoclonal antibodies and 4 cocktails, against the R346-mutated BA.4/BA.5 sublineages (Fig. 1E) ^7–14^. Cilgavimab was completely escaped by BA.4/BA.5 with R346I/T/S, resulting in the complete loss of efficacy of Evusheld (Tixagevimab+Cilgavimab) against BA.4.6, BA.4.7, and BA.5.9. The neutralizing activity of REGEN-COV (Casirivimab+Imdevimab) was also reduced due to the decreased reactivity of Imdevimab against R346-mutated sublineages. The potency of Sotrovimab is further reduced. Notably, Bebtelovimab remains highly potent and is the only surviving NAb drug approved by the U.S. food and drug administration (FDA).

Together, our findings suggest that the significant humoral immune evasion, especially against convalescents from BA.4/BA.5 breakthrough infection, contributes to the emergence and rapid spread of multiple R346-mutated BA.4/BA.5 sublineages. The decreased neutralization titers of the plasma samples from BA. 5 breakthrough-infection convalescents indicate worrisome potential reinfection of BA.4.6 after the recovery from BA.4/BA.5, and close monitoring of those subvariants should be performed.

## Acknowledgments

We thank all volunteers for providing the blood samples. This project is financially supported by the Ministry of Science and Technology of China and Changping Laboratory under the project number (CPL-1233).

## Author contributions

X.S.X and Y.C. designed the study. F.J. and Y.C. wrote the manuscript. Y.Y. and Youchun W. constructed the VSV-based pseudovirus. L.Y., Y.G., N.Z., Y.W. and F.S. performed the pseudovirus neutralization assays. F.J., W.S., A.Y. and Y.C. analyzed the neutralization and sequence data. X.H., Y. X and R.J recruited the SARS-CoV-2 vaccinees and convalescents.

## Methods

### Neutralizing antibody expression

Heavy chain and light chain amino acid sequences of the following antibodies were downloaded from the databases: LY-CoV016 (PDB: 7C01), LY-CoV555 (PDB: 7KMG), LY-CoV1404 (PDB: 7MMO), REGN10933 (PDB: 6XDG), REGN10987 (PDB: 6XDG), COV2-2196 (PDB: 7L7E), COV2-2130 (PDB: 7L7E), S309 (PDB: 6WPS), and ADG-2 (MZ439266/MZ439267). These antibodies were expressed using HEK293F cell lines with codon-optimized cDNA and human IgG1 constant regions in house. BRII-196 and BRII-198 were purchased from the manufacturer (Brii Biosciences).

### Plasma isolation

Blood samples were obtained from SARS-CoV-2 vaccinated and convalescent individuals who had been infected with BA.1, BA.2, and BA.5. Written informed consent was obtained from each participant in accordance with the Declaration of Helsinki. Whole blood samples were diluted 1:1 with PBS+2% FBS and then subjected to Ficoll (Cytiva, 17-1440-03) gradient centrifugation. After centrifugation, plasma was collected from the upper layer. Plasma samples were aliquoted and stored at −20 °C or less and were heat-inactivated before experiments.

### Pseudovirus construction and neutralization assay

SARS-CoV-2 variants Spike pseudovirus was prepared based on a vesicular stomatitis virus (VSV) pseudovirus packaging system. D614G spike (GenBank: MN908947 +D614G), BA.4/BA.5 spike (T19I, L24S, del25-27, del69-70, G142D, V213G, G339D, S371F, S373P, S375F, T376A, D405N, R408S, K417N, N440K, L452R, S477N, T478K, E484A, F486V, Q498R, N501Y, Y505H, D614G, H655Y, N679K, P681H, N764K, D796Y, Q954H, N969K), BA.4.6/BF.7 (BA.4/5+R346T) spike, BA.5.9 (BA.4/5+R346I) spike, BA.4.7 (BA.4/5+R346S) spike plasmid is constructed into pcDNA3.1 vector. G*ΔG-VSV virus (VSV G pseudotyped virus, Kerafast) and spike protein plasmid were transfected to 293T cells (American Type Culture Collection [ATCC], CRL-3216). After culture, the pseudovirus in the supernatant was harvested, filtered, aliquoted, and frozen at −80°C for further use.

Huh-7 cell line (Japanese Collection of Research Bioresources [JCRB], 0403) was used in pseudovirus neutralization assays. Plasma samples or antibodies were serially diluted in culture media and mixed with pseudovirus, and incubated for 1 h in a 37°C incubator with 5% CO_2_.

Digested Huh-7 cells were seeded in the antibody-virus mixture. After one day of culture in the incubator, the supernatant was discarded. D-luciferin reagent (PerkinElmer, 6066769) was added to the plates and incubated in darkness for 2 min, and then cell lysis was transferred to the detection plates. The luminescence value was detected with a microplate spectrophotometer (PerkinElmer, HH3400). IC50 was calculated by fitting a four-parameter logistic regression model.

---

**Figure S1.**
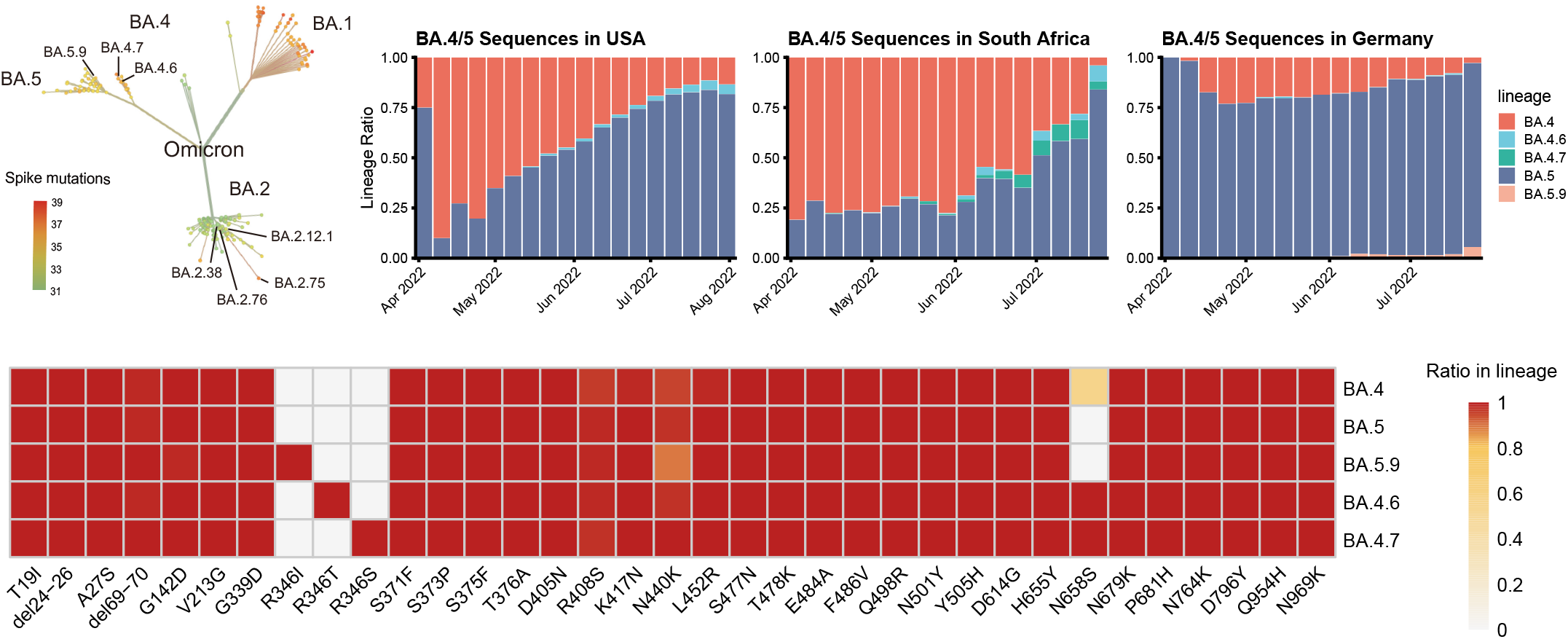
Sequence properties and growth advantage of R346-mutated BA.4/BA.5 subvariants.

**Table S1. Summarized information of the vaccinees and convalescents involved in this study.**

